# Targeting Ovarian Cancer Stem Cells by Dual Inhibition of the Long Noncoding RNA HOTAIR and Lysine Methyltransferase EZH2

**DOI:** 10.1101/2023.06.26.546524

**Authors:** Weini Wang, Yanchi Zhou, Ji Wang, Shu Zhang, Ali Ozes, Hongyu Gao, Fang Fang, Yue Wang, Xiaona Chu, Yunlong Liu, Jun Wan, Anirban Mitra, Heather M. O’Hagan, Kenneth P. Nephew

## Abstract

Persistence of cancer stem cells (CSC) is believed to contribute to resistance to platinum-based chemotherapy and disease relapse in ovarian cancer (OC), the fifth leading cause of cancer-related death among US women. HOXC transcript antisense RNA (HOTAIR) is a long noncoding RNA (lncRNA) overexpressed in high-grade serous OC (HGSOC) and linked to chemoresistance. However, HOTAIR impacts chromatin dynamics in OCSC and how this oncogenic lncRNA contributes to drug resistant disease are incompletely understood. Here we generated HOTAIR knock-out (KO) HGSOC cell lines using paired CRISPR guide RNA design to investigate the function of HOTAIR. We show that loss of HOTAIR function re-sensitized OC cells to platinum treatment and decreased the population of OCSC. Furthermore, HOTAIR KO inhibited the development of stemness-related phenotypes, including spheroid formation ability, as well as expression of key stemness-associated genes ALDH1A1, Notch3, Sox9, and PROM1. HOTAIR KO altered both the cellular transcriptome and chromatin accessibility landscape of multiple oncogenic-associated genes and pathways, including the NF-kB pathway. HOTAIR functions as an oncogene by recruiting enhancer of zeste 2 (EZH2) to catalyze H3K27 tri-methylation to suppress downstream tumor suppressor genes, and it was of interest to inhibit both HOTAIR and EZH2. In vivo, combining a HOTAIR inhibitor with an EZH2 inhibitor and platinum chemotherapy decreased tumor formation and increased survival. These results suggest a key role for HOTAIR in OCSC and malignant potential. Targeting HOTAIR in combination with epigenetic therapies may represent a therapeutic strategy to ameliorate OC progression and resistance to platinum-based chemotherapy.

## Introduction

Epithelial ovarian cancer (OC) is the fifth leading cause of cancer-related death among women worldwide. In the US, it is estimated 19,710 women will be diagnosed with OC and 13,270 women will die from OC in 2023 (1). Disease recurrence is a main contributor to the low (47.4%) overall 5-year survival rate for OC, which has only marginally improved since 1980 (2). In this regard, high-grade serous OC (HGSOC) is a highly chemoresponsive tumor with very high initial response rates to standard therapy consisting of platinum and paclitaxel. However, most women (approx. 80%) eventually develop recurrence, which rapidly evolves into chemo-resistant disease. Recurrent OC is essentially incurable.

Although chemotherapy initially decreases tumor bulk, it leaves behind residual OC stem cells (OCSCs) capable of regenerating tumors. Persistence of OCSCs at the end of chemotherapy is a critical contributor to the emergence of recurrent tumors (3,4). CSCs are characterized as being highly tumorigenic, processing the ability to self-renew, differentiate, maintain tumor heterogeneity and contribute to therapy resistance (5). OCSC maintenance requires reprogramming of the epigenome, including remodeling of chromatin modifications (6). Epigenetic regulators including long noncoding RNAs (lncRNAs) play an important role in maintaining CSC characteristics. HOX transcript antisense RNA (HOTAIR) is an oncogenic lncRNA and elevated expression of HOTAIR has been associated with many different types of malignancies and shown to promote cancer stemness and chemoresistance (7). In OC, overexpression of HOTAIR correlated with disease recurrence and poor prognosis (8, 9), and HOTAIR has been implicated as a key epigenetic regulator of CSCs by us and others (10, 11). Furthermore, we demonstrated that aberrant expression of HOTAIR was associated with the acquisition of stem cell characteristics in OCSCs (12, 13), but details of the underlying functional and regulatory mechanisms are unclear.

HOTAIR reprograms global chromatin state via recruitment of critical protein partners to its target genes. HOTAIR acts as a molecular scaffold that guides polycomb repressive complex 2 (PRC2) and lysine demethylase 1 (LSD1) complexes to chromatin, marking genes for transcriptional repression via trimethylation of histone H3 Lys 27 (H3K27me3) and H3K4 demethylation, respectively, resulting in global alterations in gene expression levels and repression of tumor suppressors (13,14. 15). By using a peptide nucleic acid (PNA) hybrid designed to block HOTAIR binding to EZH2, a core component of PRC2, we demonstrated that chemoresistant ovarian tumors were re-sensitized to platinum (13). We expanded this concept of disrupting HOTAIR-EZH2 interactions as a means of reducing the OCSC population, supporting the possibility that targeting HOTAIR and blocking its function alone or in combination with other epigenetic inhibitors (12) could potentially eliminate OCSCs and overcome OC chemoresistance.

In this study, to examine the mechanism by which HOTAIR regulates OCSCs, we generated HOTAIR knockout (KO) models using a paired gRNA targeting strategy. We demonstrated that HOTAIR KO significantly reduced the percentage of OCSCs through inhibiting EZH2-induced H3K27 trimethylation. We found alterations in both the transcriptome and chromatin accessibility of OCSCs after HOTAIR KO, as well as altered stemness associated genes and canonical NF- κB pathway-induced ALDH1A1 transcriptional activation. To develop new therapeutic strategies targeting OCSCs, we combined blocking the HOTAIR-EZH2 interaction using a PNA hybrid with an EZH2 inhibitor to block global H3K27me3 of tumor suppressors. This strategy synergistically increased chemosensitivity in vitro and in vivo. Taken together, we established that targeting HOTAIR-EZH2 could be a novel therapeutic approach to eliminate OCSCs and overcome OC recurrence.

## Material and Methods

### Cell culture and drug treatments

HGSOC cell lines Kuramochi and OVSAHO were obtained through MTA from the Japanese collection of Research Bioresources Cell Bank (JCRB). The JCRB cells were cultured in RPMI 1640 medium with 10% FBS. HGSOC cell line OVCAR3 was obtained from ATCC, and epithelial OC cell line A2780R5 (cisplatin resistance derivative of A2780) was previously generated by us (16, 17). Cell lines were maintained in culture as described previously (12, 16). Cell lines were tested for mycoplasma contamination (Lonza, cat #LT07-318) every 6 months. Cells were authenticated by short tandem repeat (S7TR) analysis in 2017 (IDEXX BioAnalytics). Cells were treated with the EZH2i (GSK503, GSK Inc.) for 72 hours followed by functional assays. HOTAIR overexpression in HOTAIR knockout cells was performed by transfecting the pAV5S vector containing a 98-mer aptamer sequence and full-length HOTAIR as we have previously described (18), empty vector was transfected as control. The aptamer was used as control for any potential RNA-dependent signaling effects.

### pspCas9(BB)-2A-GFP(PX458)-sgHOTAIR plasmid construction

The sgRNA design was based on the CHOPCHOP online CRISPR-Cas9 design tool. The sgRNA oligos targeting the different genomic regions of HOTAIR gene were synthesized and cloned into the BbsI site of PX548 (Addgene, Cat. No: 48138) construct following manufacturer’s instructions. The plasmids were transformed into DH5-alpha competent cells and sequenced using U6 promoter targeting primer by Eurofins Scientific. Effective sgRNA containing constructs were selected by T7E1 (NEB) cleavage assay following manufacturer’s instructions.

### CRISPR/Cas9

For HOTAIR KO, OC cells were co-transfected with two PX458 plasmids each containing HOTAIR single guide RNA (sgRNA1 sequence 5’-3’: TCAGGTCCCTAATATCCCGG, sgRNA2 sequence: 5’-3’: CCCGGCTTCTCATCGCTAG) using Lipofectamine 3000 according to manufacturer’s protocol. Control PX458 plasmid contained a non-targeting 20nt scramble gRNA. For RelA KO, OC cells were transfected with lentivirus based CRISPR plasmid (MISSION® gRNA, HSPD0000035812, Sigma Aldrich) targeting human RelA gene (sgRNA sequence: CTACGACCTGAATGCTGTG). CRISPR-Lenti non-targeted control plasmid (CRISPR12-1, Sigma Aldrich) was transfected into cells to serve as a control. GFP-positive single cells were isolated and sorted into 96-well plates by FACS Aria cell sorter (BD Biosciences). Cells were maintained in 96-well plates until they reached confluence and then expanded into larger plates for further analysis. Knockout status was validated by PCR, qRT-PCR (HOTAIR Forward primer: CAGTGGGGAACTCTGACTCG, HOTAIR Reverse primer: GTGCCTGGTGCTCTCTTACC), and western blotting.

### T7 Endonuclease I assay

Viable clonal cell expansions were visualized before being subjected to the following assay: 500,000 GFP+ cells of each clonal cells were used in T7EI assay for positive KO cell detection. T7 endonuclease I enzyme was purchased (New England BioLabs) and assay was performed followed by the manufacturer’s protocol.

### RNA- and ATAC-sequencing (seq) analysis

Three positive Kuramochi HOTAIR KO clonal cells and three control Kuramochi clonal cells were lysed with RLT, and RNA was extracted with RNeasy Mini kit (Qiagen) according to the manufacturer’s protocol. RNA-seq was performed as we have previously described (19). RNA-seq reads were firstly assessed using FastQC (v.0.11.5 Babraham Bioinformatics, Cambridge, UK) and trimmed by Trim Galore (v.0.6.5 https://github.com/FelixKrueger/TrimGalore) for quality control. The sequence data were then mapped to an reference genome hg38 using the Hisat2 (v.2.5) (20) and using the Samtools (v.1.15) (21, 22) to only keep uniquely mapped reads. Uniquely mapped reads were used to quantify the gene expression employing featureCounts (subread v.1.5.1) (23). The data was normalized using FPKM (fragments per kilobase of exon per million mapped fragments). Differential gene analysis was performed, and p-values were adjusted utilizing DESeq2 (v.3.14) (24).

For ATAC-seq, the resulting libraries from triplicate monoclonal cells (described above in the CRISPR/Cas9 section) were sequenced on Illumina NovaSeq 6000 with v1.5 reagent kit and paired-end 50 bp reads were generated. Illumina adapter sequences and low-quality base calls were trimmed off the paired end reads with Trim Galore (v.0.6.5 https://github.com/FelixKrueger/TrimGalore). The resulting high-quality reads were aligned to the human reference genome hg38 using bowtie2 (v.2.4.2) (25) with parameters “-X 2000 --no-mixed --no-discordant”. Samtools (v.1.15) (21, 22) was employed to map only unique reads with proper pairs. Duplicate reads were identified using Picard (v.2.21.6 https://broadinstitute.github.io/picard/) and discarded. MACS2 (v.2.2.7.1) (26) was used to identify general peaks with “-f BAMPE --keep-dup all -B -q 0.01”. BAMscale (v.0.0.5) (27) was employed to normalize and convert all bam files into bigwig files appropriate for Integrated Genome Browser (v.2.11.9) (28) and to visualize the distribution of reads along the genome.

### Motif analysis

FindMotifsGenome.pl from HOMER (v.4.11, http://homer.ucsd.edu/homer/) was utilized to identify the motifs enriched in peaks upregulated in KO and control respectively with adjusted p-value cutoff at 0.05. NF-κB motif identified by HOMER was used to identify the corresponding peaks.

### Integrated analysis

We annotated the peaks upregulated in KO and control respectively to proximal genes using ChIPseeker (v.1.28.3) (29) and only peaks located within gene promoters (upstream or downstream 3000 bases from transcription start site) were assessed. Overlapped genes were identified in both seq datasets for KO vs control (genes upregulated in RNA-seq and peak-associated genes upregulated in ATAC-seq). Hallmark gene sets downloaded from Molecular Signature Database (MsigDB) (30) and GSVA (v.1.40.1) were utilized to infer the activity scores of pathways from Hallmark for corresponding samples. GO (Gene Ontology) gene sets downloaded from MsigDB (30) were used to find stemness-related gene sets (29). Gene sets containing the term ‘stem’ from Gene Ontology are used for further analysis and then GSVA was used to infer the activity scores of pathways.

### Spheroid formation assay

500 cells were plated in 24-well ultra-low adherent plates (Corning) in triplicate with 1ml of customized stem cell medium as previously described (31). Cells were allowed to grow for 14 days in culture prior to counting. Spheroid size and morphology were assessed using a Zeiss Axiovert 40 inverted microscope with Axio-Vision software (Carl Zeiss MicroImaging). Spheres or clusters smaller than 100 µm were not counted.

### Flow cytometry and ALDEFLUOR assay

ALDH enzymatic activity was measured using the ALDEFLUOR assay kit following the manufacturer’s instructions (Stemcell Technologies) as described previously (19, 31). The ALDH-positive population was sorted and analyzed by FACS Aria II flow cytometer (BD Biosciences) or LSRII flow cytometer (BD Biosciences). ALDH-positive cells were gated using control cells incubated with the ALDH inhibitor, diethylamino benzaldehyde (DEAB).

### Luciferase reporter assay

ALDH1A1 promoter region (1031 bp) was cloned into an empty pGL3-Luc vector. Kuramochi control, NF-κB KO cells, or HOTAIR KO cells were co-transfected with renilla luciferase plasmid and pGL3-ALDH1A1-Luc or empty vector control. Transfected Kuramochi control cells were then treated with TGF-beta (10 ng/ml; Sigma) for 2 hours before measurement to serve as a positive control for activated NF-κB activity. Renilla luciferase activity was used for normalization.

### Western blot

Proteins were extracted from cells with RIPA buffer (Thermo Fisher, cat #89900) containing Pierce^TM^ protease inhibitor cocktail. Protein concentrations were quantified with the Bradford assay (Bio-Rad, #5000001) according to the manufacturer’s protocol. Lysates were subjected to SDS-PAGE and transferred to a PVDF membrane by standard methods following manufacturer’s instruction. Western blots were probed with primary antibodies p65 (Cell Signaling Technologies, cat #8242), p-p65 (Cell Signaling Technologies, cat #3033), I-κB alpha (Cell Signaling Technologies, cat #4176), histone H3 (Cell Signaling Technologies, cat #4499), GAPDH (Cell Signaling Technologies, cat #5174), and ALDH1A1 (BD Transduction Laboratories, cat# 611194). After incubation with the corresponding HRP-conjugated secondary antibodies (1:5000, Cell Signaling Technologies, cat #7074), chemiluminescent substrates from ECL kit (Thermo Scientific, cat #32106) was utilized to visualize the protein bands.

### qRT-PCR

RNA was isolated from cultured cells using the RNeasy mini kit (Qiagen) following the manufacturer’s protocol. Concentrations were determined by Nanodrop. Total RNA (2µg) was reverse transcribed with the following manufacturer’s instructions from Promega. qPCR was performed using Lightcycler 480 SYBR Green I Master kit (Roche Diagnostics) using indicated primers (IDT). mRNA expression levels were determined using Lightcycler software version 3.5 (Roche Applied Science) and normalized to EEF1A1. ddCt methods was used to determine relative mRNA levels in between groups. Primer sequences for qPCR can be found in the Supplemental Table 1 and our previous publication (12).

### Immunofluorescence

OC cells and spheroids were fixed, permeabilized and stained as described (31). Staining with ALDH1A1 antibody (1:500, BD biosciences) was followed by Alexa fluor-488 anti-mouse antibody (1:1000, Thermo Fisher). Staining with p65 antibody (1:500, Santa Cruz) was followed by Alexa fluor-568 anti-mouse antibody (1:1000, Thermo Fisher). Nuclei in spheroids were stained by Hoechst (Invitrogen). At least 10 random images from 3 independent experiments were taken using a light microscope.

### Combination index and synergism

1000 cells were plated in 96-well plates and treated with cisplatin at IC_50_ dosage for 3 hours. Cells were then treated with different single, or combination treatments as follows: GSK503: 250nM, 500nM, 750nM for 72 hours; PNA3 100nM and 200nM for 24 hours. Following treatments, the percent survival subtracted from 100% was indicative of the fraction affected (FA). Subsequent combination indices and synergism determination were calculated by the Chou-Talalay method (31).

### Mouse xenograft experiments

All animal studies were conducted based on ethical guidelines approved by the Institutional Animal Care and Use Committee of Indiana University. Kuramochi cells (1 x 10^6^) were mixed with matrigel at a 1:1 ratio and injected subcutaneously into the right flank of NSG female mice.

Tumor size was measured twice a week with a caliper, and tumor volume was determined using the formula V=½ × L × W^2^, where L is the longest tumor diameter and W is the perpendicular tumor diameter. The mice were randomized into 8 groups (n=5 or 6 mice per group) and when the tumor size reached 200 mm^3^, treatments as described were initiated (administered by intraperitoneal (IP) injection). Mice were sacrificed and tumors were collected either when the tumor volume reached 2000 mm^3^ (protocol limit) or at the end of the study. Tumors were dissociated into single-cell suspension using tumor dissociation kit (Miltenyi Biotec) in combination with the gentleMACS™ Dissociator, prior to FACS sorting and qPCR. Mouse survival in each group was analyzed by Prism 9.

### Statistical analysis

All data are presented as mean values ± SD of at least three biological experiments unless otherwise indicated. The estimated variation within each group was similar therefore Student’s *t*- test was used to statistically analyze the significant difference among different groups.

RNA-seq and ATAC-seq results are available for download at Gene Expression Omnibus (GEO) data repository at the National Center for Biotechnology Information (NCBI) under the accession number GSE144948.

## Results

### Generation of HOTAIR knockout OC cell lines

To better understand the functions of HOTAIR in cancer stemness and platinum resistance, HOTAIR knockout (KO) cell lines were generated using CRISPR/Cas9 technology. To delete the genomic region of HOTAIR, two gRNAs were used, one targeting exon1 and the other exon 6 (Suppl. Fig. S1A), resulting in complete loss of lncRNA expression through RT-qPCR in clonal populations of HGSOC cell lines Kuramochi (Fig. 1A), OVCAR3 (Suppl. Fig. S1B, C) and epithelial OC cell line A2780R5 (Suppl. Fig. S2C). To confirm that HOTAIR was deleted in the HGSOC cells line, three pairs of PCR primers expanding the genomic truncation region were used (Suppl. Figs. S2A, Kuramochi; S1B, OVCAR3), and Sanger-sequencing was used to confirm genomic truncation in all the OC KO clones (data not shown). Furthermore, as HOTAIR is a type of anti-sense lncRNA residing within the protein-coding gene HOXC11 (transcribed in the opposite direction of HOXC11), it was important to rule out any effect of depleting the HOTAIR gene on HOXC11 gene expression. We verified through RT-qPCR that HOXC11 mRNA expression in the clones from all three cell lines and clones selected for subsequent experiments was unchanged (Fig.1A, Suppl. Fig S1C, S2C).

**Figure 1.**
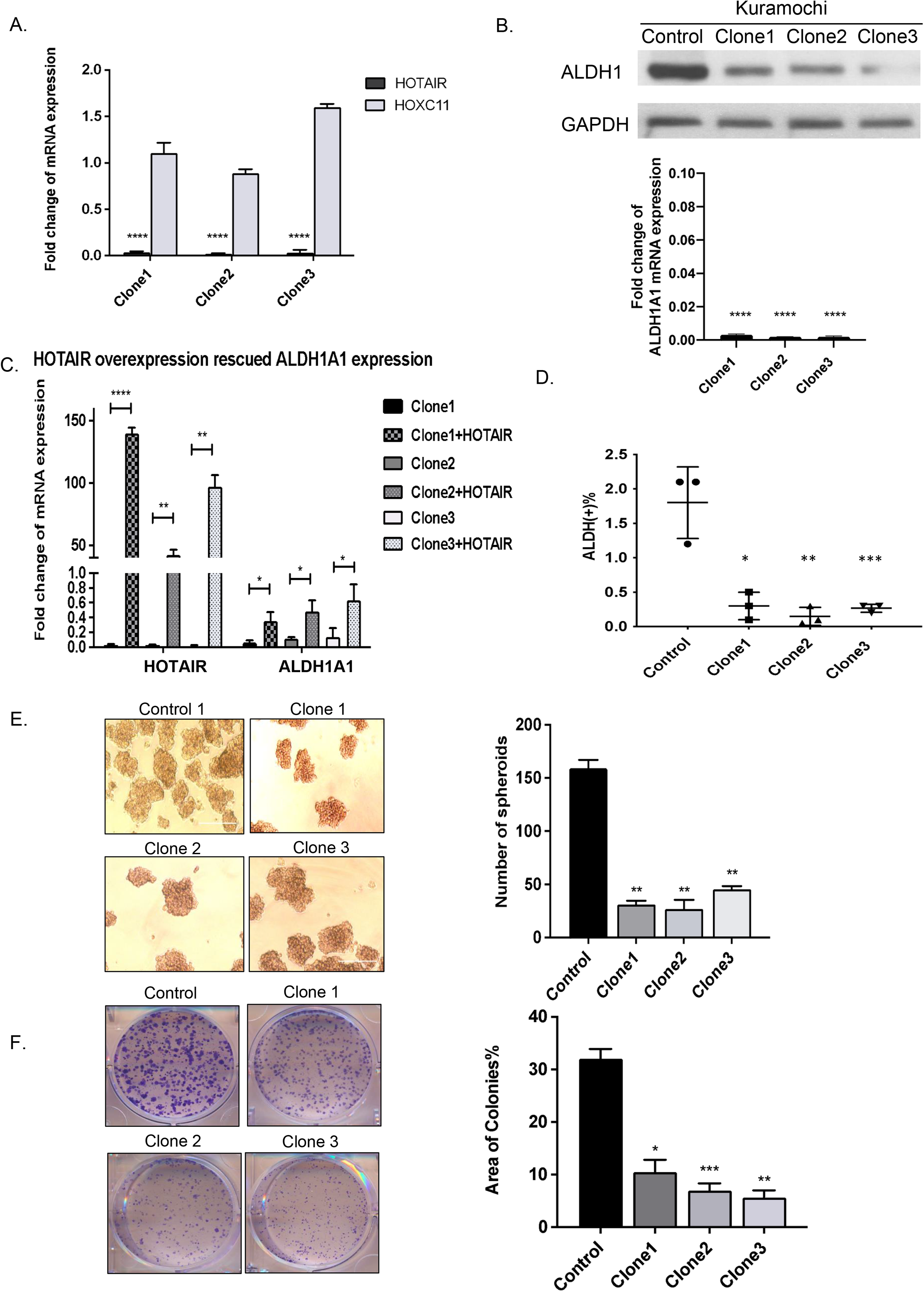
CRISPR/Cas9-mediated knockout of HOTAIR decreased ovarian cancer stemness. **A**. Expression of HOTAIR and HOXC11 genes were determined using quantitative real-time PCR (qPCR) in Kuramochi HOTAIR knockout (KO) clones versus vector control cells. **B.** (Upper) Western blot analysis of ALDH1 in Kuramochi KO versus vector control cells. GAPDH was used as loading control. (Lower) Expression of ALDH1A1 was determined by qPCR analysis in Kuramochi KO clone cells versus vector control cells. **C.** HOTAIR overexpression rescued ALDH1A1 expression in KO clones. Expression of HOTAIR and ALDH1A1 were determined by qPCR analysis. **D.** Quantification of flow cytometry showing the percent ALDH (+) cells in Kuramochi KO cells and vector control cells. **E.** Representative images of spheroid formation in 500 Kuramochi HOTAIR KO versus vector control cells. Quantification of spheroids is shown. **F.** Representative images of colony formation in 500 Kuramochi HOTAIR KO versus vector control cells. Quantification of colonies is shown. Representative data of at least three replicates. Scale bars, 100 μm. *, *P <0.05; **, P<0.01; ***, P<0.001; ****, P<0.0001*.

### HOTAIR knockout decreases OC cell stemness

A robust functional CSC marker is ALDH1A1 (16, 31). In the HOTAIR KO clones, expression of ALDH1 protein (Fig. 1B upper, Kuramochi; OVCAR3, Suppl. Fig. S1D) and ALDH1A1 mRNA (Fig. 1B lower, Kuramochi; OVCAR3 and A2780R5, Suppl. Fig. S1C and S2C, respectively) were decreased. In addition, HOTAIR KO decreased the cisplatin IC50 from 12.1 μM to 5.7 μm compared to control cells (Suppl. Fig. S2B, Kuramochi). Re-overexpression of HOTAIR in the HOTAIR KO clones rescued the ALDH1A1 mRNA expression compared to the KO cells (Fig. 1C), indicating HOTAIR is an upstream regulator of ALDH1A1. Consistent with the decreased expression of ALDH1A1, the ALDH (+) cell population, determined by ALDEFLUOR assay, significantly decreased from 1.8% to less than 0.5% in the HOTAIR KO clones (Fig. 1D, Kuramochi; Suppl. Fig. S2D, A2780R5), and immunofluorescent staining showed reduced ALDH1A1 staining in the HOTAIR KO clones compared to the control cells (Suppl. Fig S2E).

We then examined spheroid formation, a functional assay of stemness, and clonogenic activity, a sensitive indicator of undifferentiated CSCs. The ability of HOTAIR KO cells to form spheroids in ultra-low attachment plates was significantly impaired compared to controls (Fig. 1E, Kuramochi; OVCAR3 and A2780R5, Suppl. Figs. S1F and S2F, respectively). In addition, colony formation ability by HOTAIR KO clones was significantly reduced compared to control cells (Fig. 1F, Kuramochi; OVCAR3 and A2780R5, Suppl. Figs.S1G and S2G, respectively), demonstrating impaired clonal expansion ability. These results demonstrate that HOTAIR plays a key role in the cancer stemness phenotype in OC.

### HOTAIR and EZH2 regulate gene expression and genome-wide chromatin accessibility in OCSCs

HOTAIR acts as a molecular scaffold to recruit chromatin-modifying enzymes to the genome, including EZH2, and has been shown to direct EZH2 to specific gene loci in breast cancer (33). We next examined whether HOTAIR functions through EZH2 in regulating OCSC phenotypes. Control and HOTAIR KO Kuramochi cells were treated with EZH2i (GSK503), and spheroid and colony formation abilities were examined. GSK503 reduced both the number of spheroids (Fig. 2A) and colonies (Fig. 2B) formed by control HGSOC cells; however, inhibiting EZH2 in HOTAIR KO cells did not further reduce the ability of cells to form spheroids (Fig. 2A) or colonies (Fig. 2B), indicating that HOTAIR and EZH2 function in the same pathway in regulating OCSC phenotypes.

**Figure 2.**
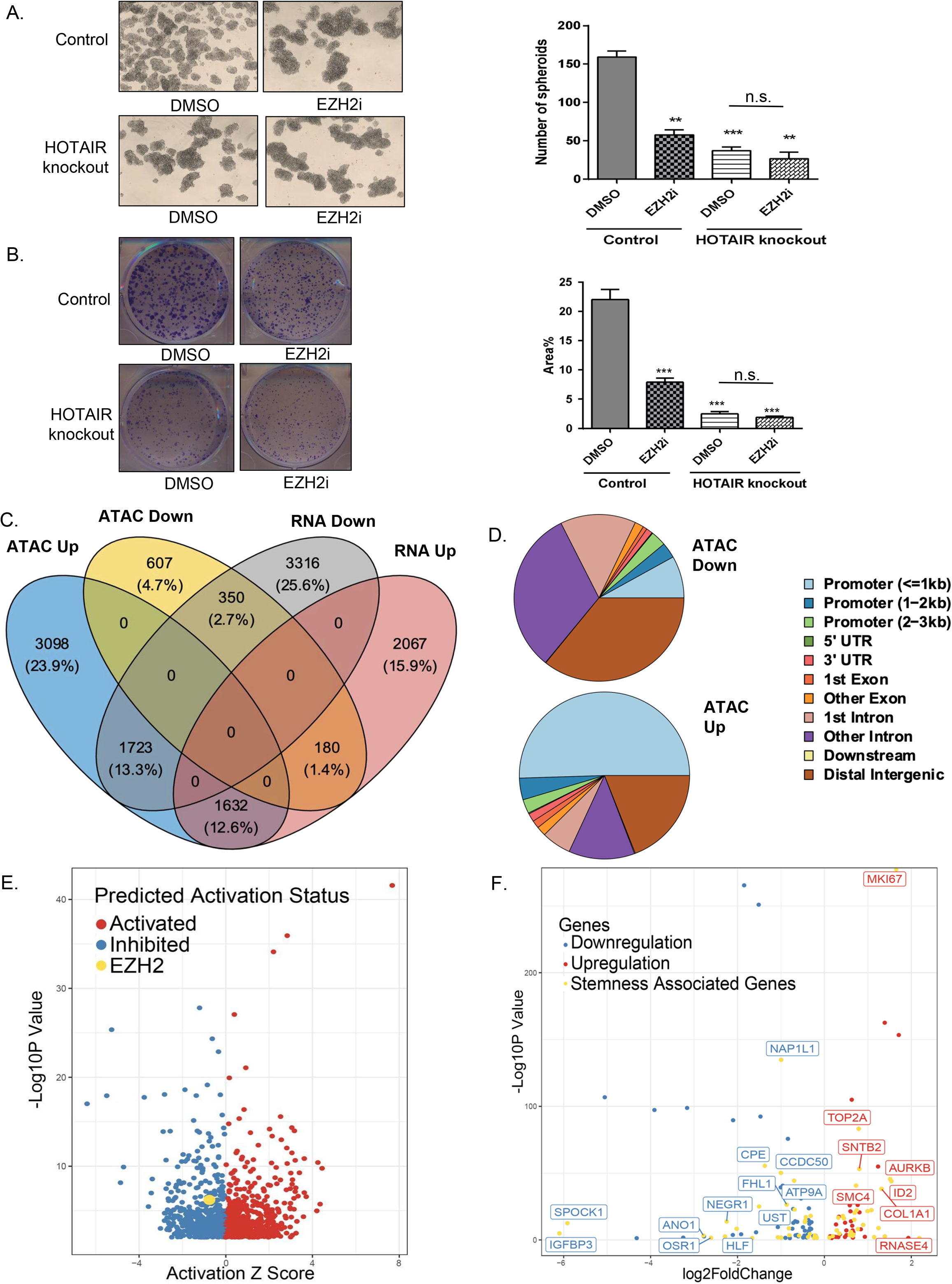
HOTAIR functions through EZH2 in regulating OCSCs. **A.** Representative images of spheroid formation (500 Kuramochi HOTAIR KO versus vector control cells were treated with GSK503 (EZH2i, 500 nM) or vehicle control (DMSO)). Quantification is shown to the right. **B.** Representative images of colony formation (500 Kuramochi HOTAIR KO versus vector control cells were treated with GSK503 or vehicle control). Quantification is shown to the right. **C.** Venn Diagram of overlap of differential peaks-associated genes in ATAC-Seq and differentially expressed genes in RNA-Seq. **D.** Pie charts for annotations of distribution of upregulated and downregulated peaks in HOTAIR KO vs Control (ATAC-Seq up/down regulated genomic regions in HOTAIR KO cells in genomic regions). Peaks were annotated based on the most proximal genes. Distribution of peaks enriched in annotated genomic regions is shown. **E.** Volcano plot of activity z scores for upstream regulators analyzed by Integrative Pathway Analysis software. Activation z scores accompanied by p values were inferred. **F.** Volcano plot of differentially expressed EZH2 target genes with adjusted p-value <0.05. Stemness-associated genes are represented by yellow dots. *, P <0.05; **, P<0.01; ***, P<0.001; ****, P<0.0001.

Because oncogenic HOTAIR and EZH2 are known to reprogram chromatin status and alter chromatin modifications (10, 34), we performed an Assay for Transposase-Accessible Chromatin using sequencing (ATAC-seq) to assess genome-wide changes in the chromatin landscape induced by HOTAIR KO. In addition, to study the impact of HOTAIR KO on the transcriptome, we performed RNA-seq and then integrated the transcriptomic and epigenomic data. ATAC-Seq data showed HOTAIR KO led to altered chromatin accessibility (Suppl. Fig. 3A). There were in total 29,442 peaks with differential accessibility between HOTAIR KO and control, of which 14,625 peaks were up-regulated and 14,817 peaks were down-regulated. To assess whether the altered chromatin landscape was associated with differential gene expression, integrated analysis of RNA-seq and ATAC-seq was performed by comparing differentially accessible peaks in the immediate vicinity of differentially expressed genes. From the overlapping set, 1,982 genes with altered chromatin accessibility and differential expression were identified, including 350 downregulated genes and 1632 upregulated genes associated with ATAC-seq peaks (Fig. 2C). Moreover, we observed both up-regulated and down-regulated peaks to be mainly located in distal intergenic regions (Fig 2D, upper pie chart). We found that a higher percentage of up-regulated peaks occurred in promoter regions less than 1 kb in HOTAIR KO cells, indicating that HOTAIR most likely functions near promoter regions to make chromatin less accessible and down-regulate genes (Fig. 2D, lower pie chart). By analyzing the distribution of upregulated genes and downregulated peaks in HOTAIR KO versus control samples, 50.5% of the peaks associated with upregulation of gene expression resided in the promoter regions, whereas only 8.15% of peaks associated with decreased expression were in gene promoters. Moreover, by integrating chromatin accessibility and changes in gene expression, a total of 133 overlapping pathways, 40 downregulated and 93 upregulated pathways, were identified (Suppl. Fig. S3B). Among these, GO terms such as “hematopoietic stem cell differentiation” and “mesenchymal stem cell differentiation” were enriched in the downregulated gene group in HOTAIR KO cells compared with control (Suppl. Fig. S3C).

We then investigated whether gene expression changes were linked to the HOTAIR-EZH2 interaction. Integrative Pathway Analysis (IPA) of RNA-seq data revealed that EZH2 was one of the inhibited upstream regulators of HOTAIR in regulating targeted genes (Fig. 2E), and overall, 142 EZH2 target genes were identified in our dataset (Fig. 2F). By integrating the DEG genes with the stemness-related genes in the GO database, we found 59 stemness related genes were enriched (Fig. 2F). These data suggest that HOTAIR may function through EZH2 in altering chromatin landscapes and to regulate OCSCs.

### HOTAIR alters multiple stem- and cancer-associated pathways in OCSCs

Bioinformatic analysis of the transcriptomic data of HOTAIR KO and control cells identified a total of 2501 genes with altered expression (adjusted p-value<0.05, log2FC>2; volcano plot; Suppl. Fig. S3D). RNA-seq revealed a panel of essential stemness-related genes were downregulated in HOTAIR KO cells compared to control cells, including ALDH1A1, SOX4, Sox9, ZEB1, BMI1 (Fig. 3A). Furthermore, key stemness-related pathways tended (two out of three clones) to be downregulated by HOTAIR KO, including Notch, TGF-β, TNF signaling via NF-κB, Hedgehog, Wnt/β-catenin, PI3K/AKT, and KRAS signaling (Fig. 3B, Suppl. Table 2). We then applied Ingenuity Pathway Analysis (IPA) analysis and HALLMARK category of Molecular Signature Database to the DEGs regulated by HOTAIR and visualized gene expression changes between experimental groups using a hierarchical clustering heatmap. As shown in Suppl. Fig. S3C, overlapping molecular functions and predicted key biological processes consistent with tumor progression (angiogenesis, hypoxia, epithelial-mesenchymal transition, apoptosis), tumor microenvironment (metalloproteases (MMPs- MMP1,2,10,16), collagens, COL1A1, COL1A2, CXCL12) and drug resistance (drug efflux transporters, ABCB1, ABCC2, FOXO6, PI3K3) were observed.

**Figure 3.**
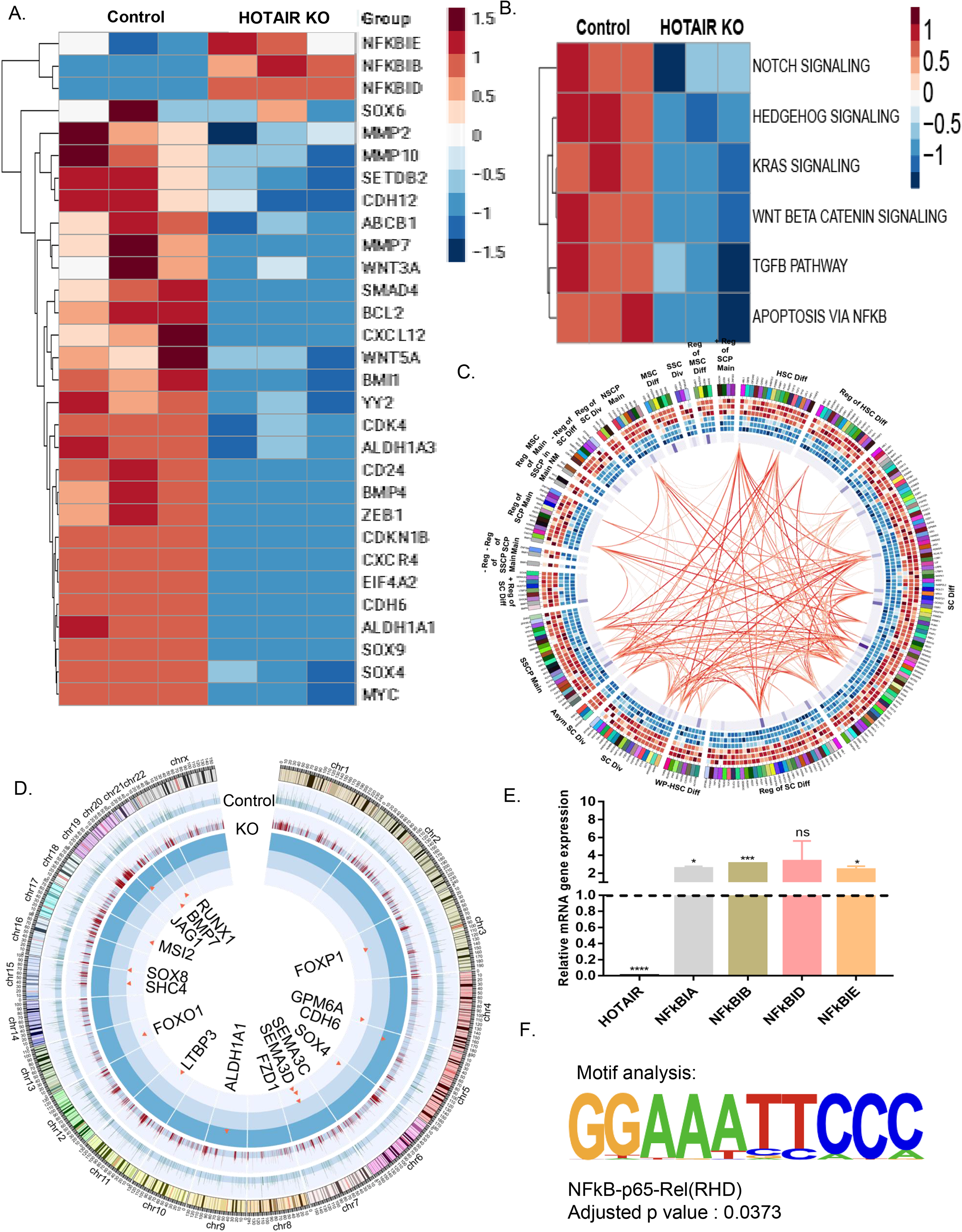
NF-κB is a target of HOTAIR-EZH2 in regulating OC stemness. **A.** Heatmap of RNA-seq data for stemness-related genes between HOTAIR KO and control Kuramochi cells. **B.** Heatmap of RNA-seq data for stemness-related signaling pathways between the two groups. **C.** CIRCOS plot for combined ATAC-Seq and stemness-related transcriptomic profiles of two groups, summarizing the differential peaks from Control and HOTAIR KO groups. The tracks from outermost to innermost represent intensity of upregulated differential peaks in Control group, upregulated differential peaks in KO group and log2foldchange of stemness-related genes upregulated in Control group. **D.** CIRCOS plot for upregulated stemness-related genes (from Gene Oncology database) in Control. The tracks from outermost to innermost represent genes (different color codes), Control group (3 tracks), HOTAIR KO group (3 tracks) and log2foldchange of genes between 2 groups; Gradient colors from blue to red indicate the gradient changes from lowest to highest expression level. Links in the center represent significant Pearson correlation coefficients (PCCs) between 2 genes. Gradient colors of red indicate the gradient changes from lowest to highest PCC. **E.** RT-qPCR analysis for NFκBIA in HOTAIR KO cells relative to control cells. **F.** Motif analysis of upregulated NFκB motif in control group. *, P <0.05; **, P<0.01; ***, P<0.001; ****, P<0.0001.

Further analysis of DEGs integrated with stemness-related gene sets within the GO database revealed decreased stemness-related genes in HOTAIR KO cells and altered stem cell biological processes, including stem cell differentiation, division, maintenance, and cell renewal (CIRCOS plot, Fig. 3C). Furthermore, CIRCOS plot showing integrated analysis of DEGs in RNA- and ATAC-seq identified a group of genes linked to stemness significantly altered by HOTAIR KO, including ALDH1A1, FOXO1, FOXP1, SOX4/8, and CDH6 (Fig. 3D). These data suggested that HOTAIR KO remodeling of chromatin resulted in repression of stemness-related genes. Altogether, these results suggest that HOTAIR regulates the expression of multiple stemness-associated genes by modifying their chromatin accessibility to promote the OCSC phenotype.

### NF**κ**B is a key HOTAIR-EZH2 target contributing to OC stemness phenotype

Previous work in OC has demonstrated an essential role for the non-canonical NF-κB pathway in OC stemness (35) and we have reported the RelA-mediated canonical NF-κB transcription factor pathway was shown to contribute to chemoresistance in OC (18). As shown in the heatmap of DEGs from control versus HOTAIR KO cells, NF-κB was one of the top upstream regulators, and a group of NF-κB inhibitory factors (NFκBIB, NFκBID, NFκBIE) was strongly upregulated in HOTAIR KO cells (Fig. 3A), and qPCR analysis demonstrated that NFκBIA was also upregulated in HOTAIR KO cells (Fig. 3E). To reveal possible functional pathways involving HOTAIR, we performed motif enrichment analysis by HOMER on the up- and down-regulated peaks, identified through ATAC-seq (Suppl. Fig. 3A). The top motifs enriched in down-regulated peaks involved several transcription factors, many of which were associated with stemness, including OCT4, SOX2, SOX10 and NANOG (Suppl. Fig. S3E). Furthermore, NF-κB-p65 was a top transcription factor in downregulated peaks in HOTAIR KO cells (Fig. 3F; p= 0.0373, Suppl. Fig. S3E). Based on these data, we carried out mechanistic and functional analyses of canonical NF-κB pathway in regulating OCSCs.

ATAC-seq track analysis showed upregulated peaks in NFκBIA promoter in the HOTAIR KO cells compared to control (Suppl. Fig.3F), demonstrating that HOTAIR KO resulted in transcriptional activation of NF-κBIA. Furthermore, immunofluorescent staining of adherent Kuramochi cells and spheroids showed that HOTAIR KO reduced (>10 fold) nuclear localization of NF-κB subunit p65 (Fig. 4A-B, Suppl Fig. S4A), and decreased ALDH1A1 staining (Fig. 4A). Analysis of cytoplasmic and nuclear protein fractions revealed significantly reduced nuclear NF- κB expression after HOTAIR KO, which correlated with reduced ALDH1 expression (Fig. 4C). Similarly, NF-κB inhibitor Bay-11 treatment reduced ALDH1 expression, however, no further reduction of ALDH1 expression was observed in the HOTAIR KO cells (Suppl. Fig. S4B). Downregulation of NF-κB target genes IL6 and IL8 was observed after treatment of Kuramochi cells with NF-κB inhibitor Bay-11 (Suppl. Fig. S4C). Bay 11 treatment also decreased spheroid formation ability (Fig 4D), NOTCH3 gene expression (Suppl. Fig. S4D) and the percent ALDH+ cells (Suppl. Fig. S4E-F) in Kuramochi cells

**Figure 4.**
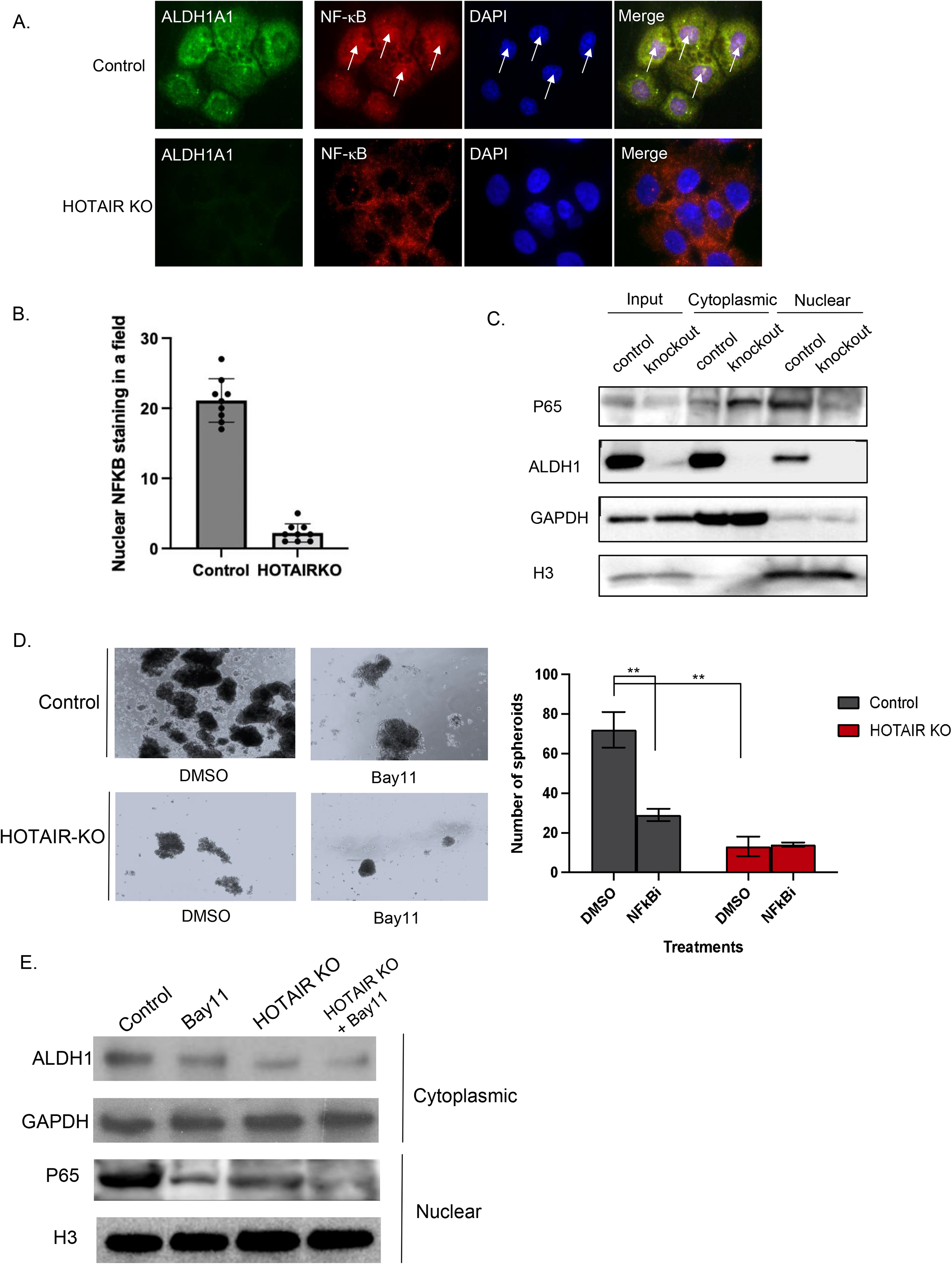
NFκB inhibition reduces OCSC level. **A.** Immunofluorescent images of ALDH1A1 (green), RelA (red) and DAPI (blue) in Kuramochi HOTAIR KO versus vector control cells. Arrows point to nuclear RelA localization. **B.** Bar graph represents the number of cells with nuclear RelA staining determined based on average of 50 cells counted at 40X magnification. **C.** Western blot of NFκB and ALDH1 in nuclear and cytoplasmic fractions from Kuramochi HOTAIR KO and vector control cells. GAPDH was used as loading control for cytoplasmic protein and H3 was used as loading control for nuclear protein. Whole cell lysate used for input. **D.** Representative images of spheroid formation in Kuramochi HOTAIR KO versus vector control cells treated with Bay-11 (7.5 μM) or vehicle control. Quantification of spheroids per image to the right. Western blot analysis of NFκB and ALDH1 in nuclear and cytoplasmic fractions from Kuramochi HOTAIR KO and vector control cells treated with Bay-11 (7.5μM, 24 hr treatment) or vehicle control. **, P<0.01.

To further test whether HOTAIR and NF-κB are functioning in the same pathway, control versus HOTAIR KO cells were treated with Bay-11 and subjected to spheroid formation assay. Reduced spheroid forming ability was observed in control cells treated with Bay-11 and in HOTAIR KO cells, but NF-κB inhibitor treatment did not further reduce spheroid formation in HOTAIR KO cells (Fig. 4D). Moreover, although decreased levels of NF-κB were observed in nuclear protein fractions of HOTAIR KO and Bay11-treated cells compared to control untreated cells (Fig. 4E), only a modest effect of Bay11 treatment on HOTAIR KO cells was observed. Collectively, these results demonstrated that HOTAIR functions through activated NF-κB in regulating OCSCs.

### NF-κB expression drives OCSC phenotype

Although NF-κB signaling was shown to be essential for regulating stemness in OC cells, the underlying mechanism remained incompletely understood. HOTAIR KO reduced the chromatin accessibility around the ALDH1A1 promoter region (Fig. 5A). To investigate whether NF-κB signaling contributed to this change in the chromatin environment, we generated NF-κB (p65 subunit) KO Kuramochi cells using CRISPR/Cas9 technology and performed western blot and RT-qPCR analysis for stemness-related genes. p65 KO reduced ALDH1 protein and ALDH1A1 mRNA expression (Fig. 5B-C), as well as NANOG and NOTCH3 expression (Fig. 5C) and NF- κB target genes (Fig. 5D) in NF-κB KO cells. NF-κB KO also decreased the ALDH+ cell population compared to control (Fig. 5E; Suppl. Fig. 4F). To determine if NF-κB regulated transcriptional activation of ALDH1A1, we performed an ALDH1A1 promoter luciferase reporter assay. HOTAIR KO decreased transactivation of ALDH1A1 reporter gene expression at a similar level as NF-κB knockout (Fig. 5F), indicating a role for HOTAIR-dependent NF-κB activity in regulating ALDH1A1 expression in OCSCs.

**Figure 5.**
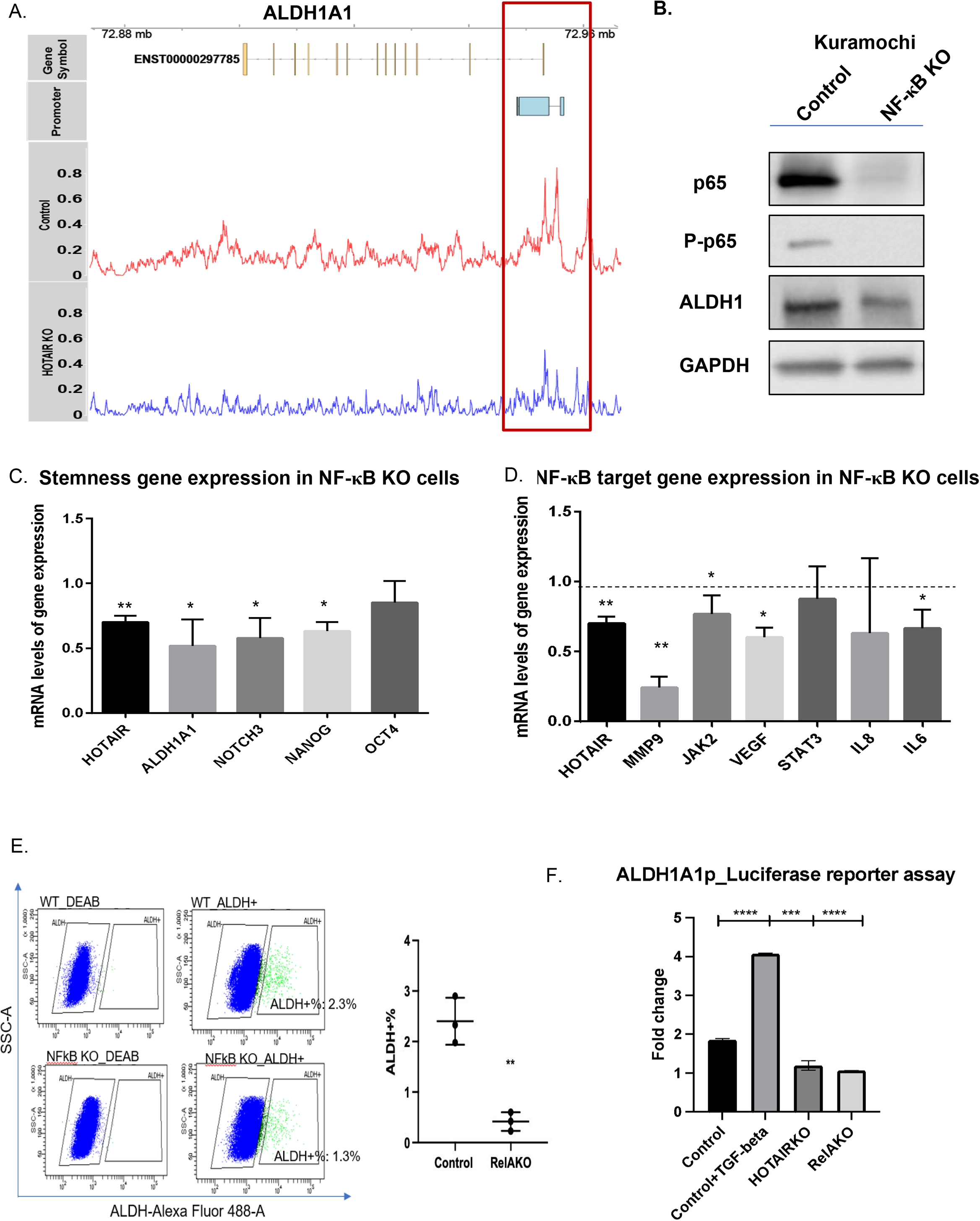
Inhibiting NFκB activity decreased OC stemness via regulating ALDH1A1 transcription. **A.** Distribution of genome coverage from different samples in ALDH1A1 genomic region and the red box represents the region with differential peaks between two groups. **B.** Western analysis of p65, p-p65 and ALDH1 protein expression in NFκB KO versus vector control cells. GAPDH was used as loading control. **C.** Expression of stemness related genes were determined by RT-qPCR analysis in NFκB KO cells versus vector control cells. **D.** Expression of NFκB target genes in NFκB KO versus vector control cells were measured by RT- qPCR. **E.** Flow scatter plot (left) and quantification of flow analysis (right) for %ALDH+ cells in Kuramochi NFκB KO versus vector control cells. **F.** OC cells were co-transfected with pGL3- ALDH1A1-Luc and renilla luciferase plasmid vector (pRL) or pGL3-Luc and pRL. Luciferase signals recorded 3 hours after drug treatment. Renilla luciferase activity used for normalization. qPCR was performed in duplicate. Average fold changes (± SD) of relative luciferase unit (RLU) compared with pGL3 are shown (*n*= 3). *, P <0.05; **, P<0.01; ***, P<0.001; ****, P<0.0001.

### Inhibiting HOTAIR and EZH2 resensitized OC cells to chemotherapy

Based on the mechanistic findings above, it was of interest to target both EZH2 and HOTAIR in OCSCs. We took a pharmacological approach and drug synergism was tested by using multiple dosages of EZH2 inhibitor GSK503 (250nM-750nM) and HOTAIR-EZH2 interaction inhibitor PNA3 (100-200nM), in combination with IC_50_ dosage of cisplatin (CDDP; 12.1μM) in Kuramochi cells. Synergistic inhibitory effects based on combination index value (CI value) lower than 1 were observed using a cell viability assay (Fig. 6A, Suppl. Fig. S5A). Based on these results, we performed an in vivo study. First, we performed a pilot drug tolerability study using NSG mice without tumor cell injection (n=6 female mice per group). Mice were treated with GSK503 (50 mg/kg, IP) 5 days per week and/or PNA3 (100 nM/kg, IP) plus cisplatin (2 mg/kg, IP) administered biweekly for four weeks. Bodyweight (BW) was monitored during the study. Throughout the observation period (29 days), BW was not different among treatment groups compared to control, indicating that the treatments were tolerated in all groups (Suppl. Fig. S5B). We then examined activity of the 50 mg/kg GSK503 dose on H3K27me3 in OC tumors. Mice were injected s.c. with 10^6^ Kuramochi cells, and GSK503 (50 mg/kg, IP) was given daily beginning when tumors reached 200mm^3^ and continued for two weeks. Tumors were harvested, and western blot analysis for H3K27me3 was performed. GSK503 effectively reduced H3K27me3 levels (Suppl. Fig. S5B), demonstrating activity at the tumor level.

**Figure 6.**
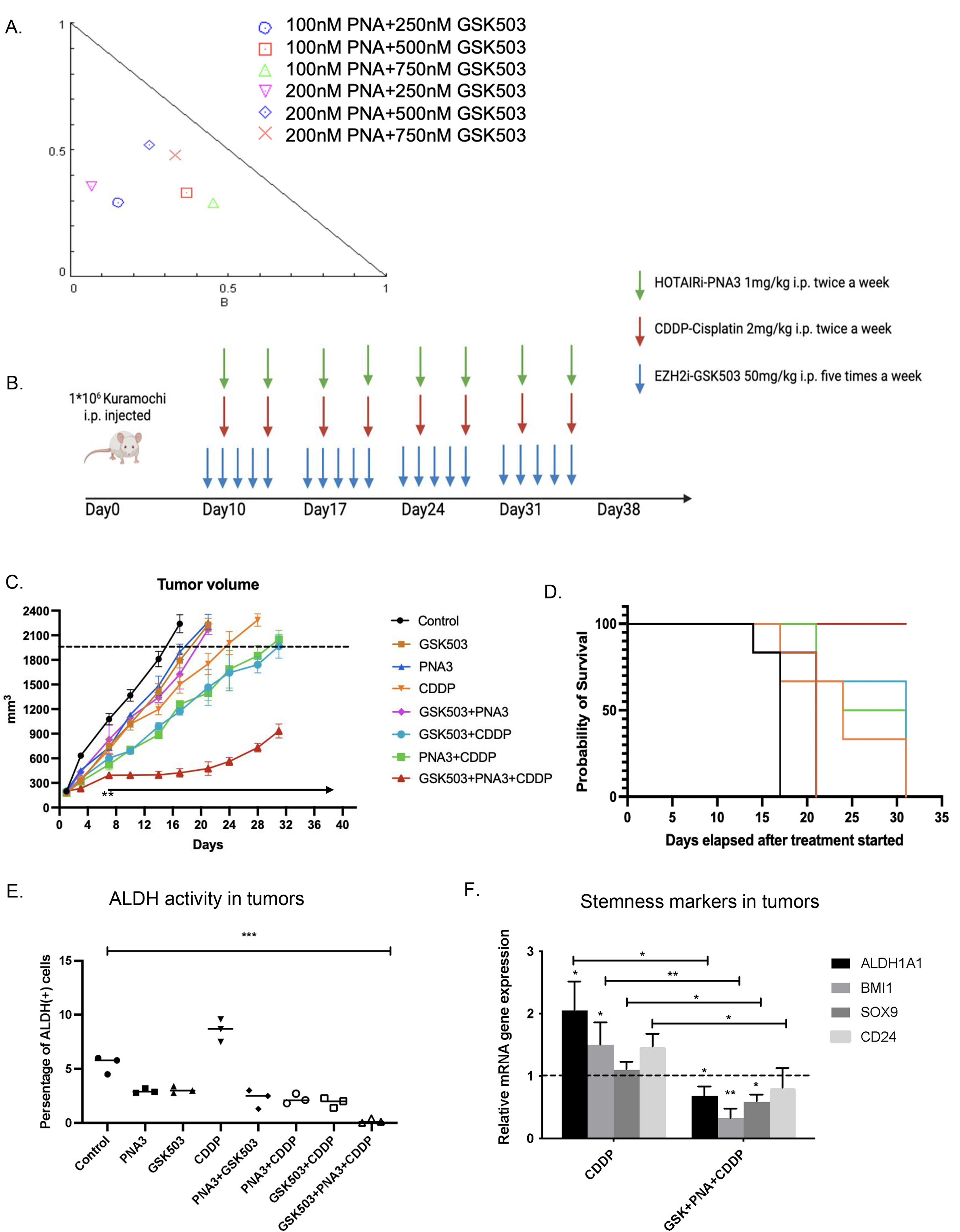
Therapeutically targeting HOTAIR and EZH2 resensitizes cells to chemotherapy *in vivo.* **A.** Synergy test of GSK503+PNA3 in Kuramochi cells treated with cisplatin at IC50 dosage (12.1 μM). Cells were first treated with GSK503 for 72hr and PNA3 for 3hrs, following with 3hr of cisplatin treatment. Effects and combination index (CI) value of different combination treatments. **B.** Scheme of *in vivo* treatment. **C.** Tumor volume of different treatments throughout the study. **D.** Survival curves for different treatments. **E.** Flow analysis of %ALDH+ cells in tumors from each treatment group determined using the ALDEFLUOR assay. **F.** Expression of different stemness-related genes were measured by RT-qPCR in cisplatin and triple combination treated tumor samples versus control tumor samples. *, P <0.05; **, P<0.01; ***, P<0.001.

Based on the results of the above pilot studies, combination treatments were performed (Fig. 6B). Kuramochi cells (1 x 10^6^) were implanted via s.c. injection and once tumor size reached 200mm^3^, mice were randomized (n=6, NSG female mice per group) and treatments were initiated. Bodyweight and tumor volume were monitored twice a week. As shown in Fig. 6C, moderate decrease in tumor volume was observed in groups treated with single drug regimens compared to control, and two-drug combinations further decreased tumor volume. However, the most significant reduction in tumor volume was observed in the group treated with GSK503+PNA3+CDDP (Fig. 6C); furthermore, the triple combination increased survival (100% triple vs. control, or double combination alone; Fig. 6D).

To determine the ALDH+ percentage at the end of the study, mice were sacrificed when tumor size reached 2000 mm^3^, tumors were collected and dissociated to single-cell suspension for FACS analysis. Cisplatin treatment significantly increased the percent ALDH+ (ALDH+%) cells from 5.5% to 8.4% (Fig. 6E). The triple combination treatment nearly eliminated the ALDH+ cells compared to either single treatment, double combination treatment or control groups (Fig. 6E, Suppl. Fig. S5D). In addition, expression of stemness-related genes (ALDH1A1, BMI1, SOX9, CD24) was examined in the harvested tumors. Cisplatin treatment increased ALDH1A1 and BMI1 expression compared to vehicle control (Fig. 6F). The triple combination significantly decreased ALDH1A1, BMI1 and SOX9 expression compared to control and compared to the cisplatin treatment group (Fig. 6F). Taken together, we concluded that inhibiting HOTAIR and EZH2 in combination with platinum-based chemotherapy can overcome chemoresistance in OC through efficiently targeting the OCSC population, decreasing tumor burden, and increasing survival.

## Discussion

In ovarian cancer, the majority of patients eventually develop recurrence after treatment with platinum-based chemotherapy. Failure to eradicate OCSCs contributes to disease relapse and drug resistance. Understanding the mechanism(s) responsible for OCSC survival and developing strategies to target these recalcitrant cells are of critical interest to the field. In the present study, we demonstrate that IncRNA HOTAIR is a key modulator of OCSCs. We found that HOTAIR functions through EZH2 to regulate downstream target genes and blocking the HOTAIR-EZH2 interaction and EZH2 activity resensitized OC cells to chemotherapy. We identified the HOTAIR-EZH2/NF-κB/ALDH1A1 axis as essential for OCSCs phenotypes.

Although chemotherapy initially decreases tumor bulk, it leaves behind residual OCSCs capable of regenerating tumors. OCSC maintenance requires reprogramming of the epigenome, including remodeling of chromatin modifications. Epigenetic drugs can be used as differentiation therapy resulting in conversion of CSCs to more differentiated cancer cells susceptible to conventional chemotherapy. Platinum resistance is linked to epigenomic alterations, including increased promoter DNA methylation and histone modifications, and we and others have investigated the potential of epigenetic therapy approaches to target these modifications in OCSCs to overcome chemoresistance (6). HOTAIR is an oncogenic IncRNA that functions as a scaffold for EZH2 in regulating downstream target gene expression (10, 35) and contributes to CSC stemness (13, 38, 39). We had previously demonstrated that HOTAIR is overexpressed in OCSCs, and overexpressing HOTAIR promotes OCSCs phenotypes (12). In this study, we used paired gRNA targeting CRSPR/Cas9 technology to KO HOTAIR specifically in OC cells and validated this approach as an effective tool to deplete expression of the IncRNA in OC cell lines rather than using traditional RNAi or shRNA techniques. We demonstrate that depletion of HOTAIR functionally decreases OCSC phenotypes and malignant potential and reprograms the cells to a less stem-like phenotype. Moreover, re-expressing HOTAIR in the HOTAIR depleted cells rescues ALDH1A1 expression, indicating that ALDH1A1 is a downstream target of HOTAIR in regulating OC stemness, but whether this is through direct regulation remains to be determined.

Here we demonstrate for the first time that combining epigenetic treatment with HOTAIR-EZH2 specific inhibitors reduces OCSCs phenotypes in vitro and in vivo, suggesting that HOTAIR and EZH2 function in the same pathway in regulating OC stemness programs. Integration of EZH2 target genes and DEGs identified by HOTAIR KO revealed a set of stemness-related genes that are co-regulated by HOTAIR-EZH2. Integration analysis of RNA-seq and ATAC-seq revealed that HOTAIR altered the global chromatin landscape of stemness-related genes, mostly through regulating EZH2 methyltransferase targeting sites. Integrated analysis further showed that ALDH1A1 expression and chromatin patterns changed significantly upon HOTAIR depletion. It is possible that this is not only due to the changes in the signaling cassettes, such as the NF-κB signaling pathway, but could also be due to the translocation of EZH2 methyltransferases from previous targeted sites to different genomic regions. An additional integrated analysis, such as ChIP-seq for EZH2 and H3K27me3 in the context of HOTAIR KO, would be necessary to examine this possibility.

As NF-κB is constitutively activated in the OCSCs, and decreased NF-κB expression inhibits ALDH1A1 transcription, we suggest that NF-κB is a direct regulator of ALDH1A1. In support of these findings, ATAC-seq revealed that the NF-κBIA promoter region is opened by HOTAIR depletion, indicating the canonical NF-κB pathway is involved in the altered OCSCs phenotypes, and we are currently investigating the role of altered NF-κB signaling in OC stemness and signaling inactivation by HOTAIR KO. Our findings agree with a recent study in breast cancer showing a functional role for HOTAIR in CSCs by a similar mechanism involving HOTAIR recruitment of PRC2 to inhibit IκBα expression and NF-κB activation (38).

We show a synergistic effect of genetic and pharmacological inhibition of both EZH2 and HOTAIR in sensitizing the OC cells to chemotherapy in vitro and in vivo. These findings suggest that HOTAIR and EZH2 co-regulate OCSCs phenotypes and are also involved in other pathways that contribute to chemoresistance. Studies have shown that HOTAIR functions through EZH2 in regulating anoikis resistance via modulating anoikis-related genes, such as ZEB1, TWIST1, and N-cadherins (39–41). Moreover, HOTAIR-EZH2 interactions were demonstrated to enhance DNA damage repair in different studies (10, 19, 43–45), suggesting that HOTAIR and EZH2 not only contribute to OCSC phenotypes, but also to other mechanisms of chemoresistance.

There is an urgent need to develop new therapeutic strategies to target OCSCs and overcome chemoresistant OC. We show that HOTAIR can promote OCSC self-renewal and drive tumor growth through NF-κB signaling-mediated activation of key stemness factors. Therapeutically, inhibiting HOTAIR and EZH2 in combination can overcome chemoresistance and improve survival by targeting the OCSC population. Targeting HOTAIR in combination with epigenetic therapies may represent a therapeutic strategy to prevent tumor relapse in OC.

## Acknowledgments

We thank Dr. Michael McCabe (GlaxoSmithKline Inc.) for providing GSK503 and helpful discussions. We thank Dr. Vaishnavi Muralikrishnan (Indiana University School of Medicine) for helpful discussions. We thank the Flow Cytometry Core Facility (Indiana University, Bloomington, IN) and Christiane Hassel for technical assistance with flow cytometry and the Light Microscopy Imaging Center (Indiana University, Bloomington, IN; Dr. Sid Shaw, Director) and Dr. Jim Powers for user training. We thank the Shared Facilities of the Indiana University Melvin and Bren Simon Comprehensive Cancer Center, the Genomics Core (Yunlong Liu, PhD, Core Director) and the Cancer Bionformatics Core (Jun Wan, PhD, Core Director). We thank Dr. Chengzu Long, Dr. Qiaoyan Yang and Dr. Ni-Huiping Son (New York University School of Medicine and Langone Health) for generously sharing the protocols and experience with CRISPR/Cas9 gene editing techniques.

## Supplemental Figure Legends

**Supplemental Figure S1.**
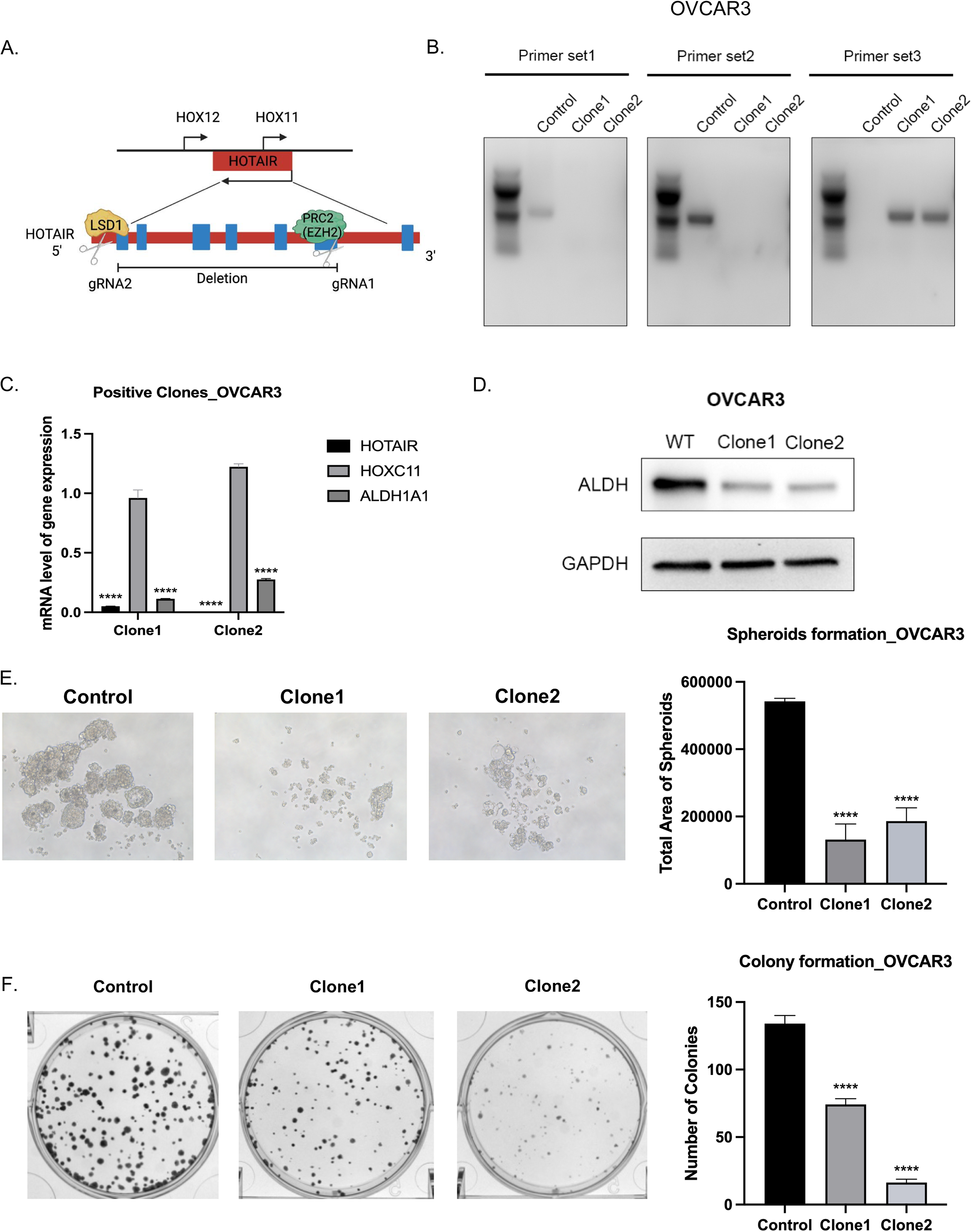
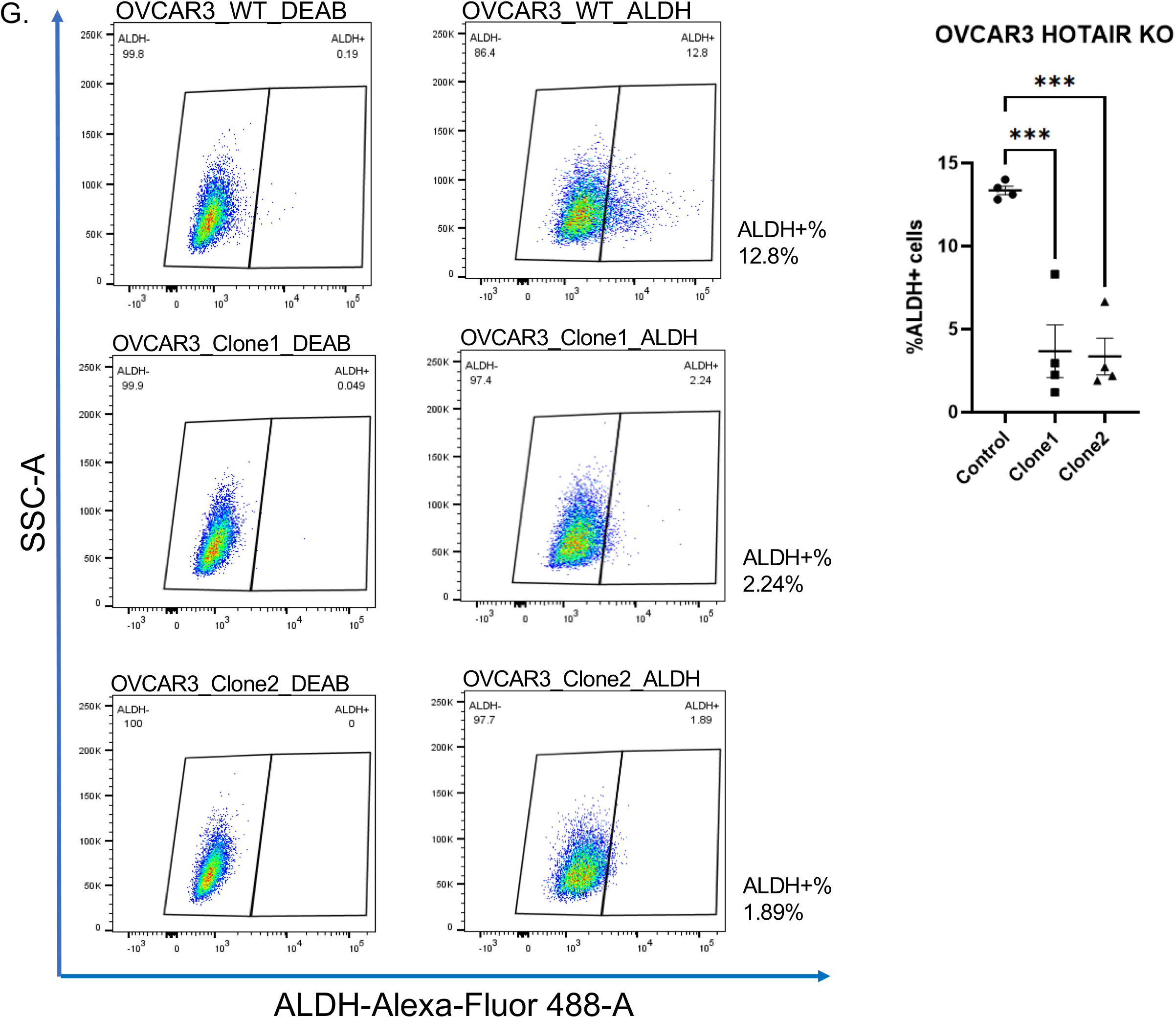
CRISPR knockout of HOTAIR decreased OC stemness. **A.** Diagram of dual gRNAs used for HOTAIR genomic deletion. Truncated region generated by dual gRNAs was measured by PCR. **C.** Expression of HOTAIR, HOXC11and ALDH1A1 genes were determined using quantitative real-time PCR in high grade serous OVCAR3 HOTAIR knockout (KO) clones versus vector control cells**. D.** Western blot analysis of ALDH1 in OVCAR3 KO versus vector control cells. GAPDH was used as loading control. **E.** Representative images of spheroid formation in OVCAR3 HOTAIR KO versus vector control cells. Quantification of spheroids is shown to the right. **F.** Representative images of colony formation in OVCAR3 HOTAIR KO versus vector control cells. Quantification of colonies is shown to the right. Representative data of at least three replicates Scale bar, 100uM*, ****, P<0.0001.* **G.** Scatter plot of the ALDEFLUOR assay showing the percent ALDH (+) cells in OVCAR3 KO cells and control cells (left). Quantification of flow cytometry (right) (*\*\*\*, P<0.001*).

**Supplemental Figure S2.**
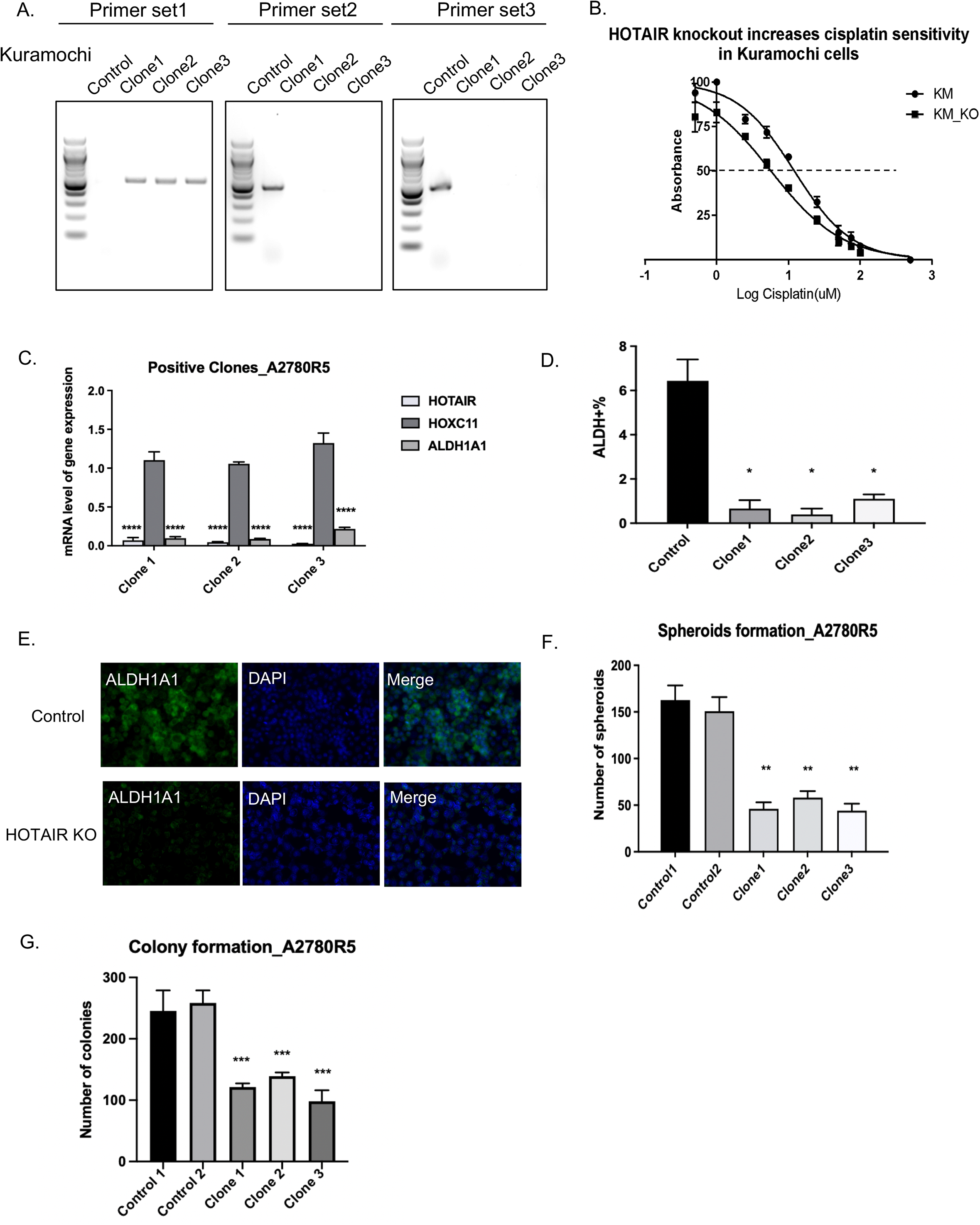
**A.** Truncational region generated by dual gRNAs was measured by PCR. **B.** IC50 of Kuramochi HOTAIR KO clonal cells versus vector control cells was determined by MTT assay. **C.** Expression of HOTAIR, HOXC11and ALDH1A1 genes were determined using quantitative real-time PCR in A2780R5 HOTAIR knockout (KO) clones versus vector control cells**. D.** Quantification of flow cytometry showing ALDH(+) cell population in A2780R5 KO clonal cells and vector control cells. **E.** Immunofluorescent staining of ALDH1A1, DAPI in A2780R5 HOTAIR KO clonal cells versus vector control cells. **F.** Quantification of spheroids in A2780R5 HOTAIR KO clonal cells versus vector control cells. **G.** Quantification of colonies in A2780R5 HOTAIR KO clonal cells versus vector control cells.

**Supplemental Figure S3.**
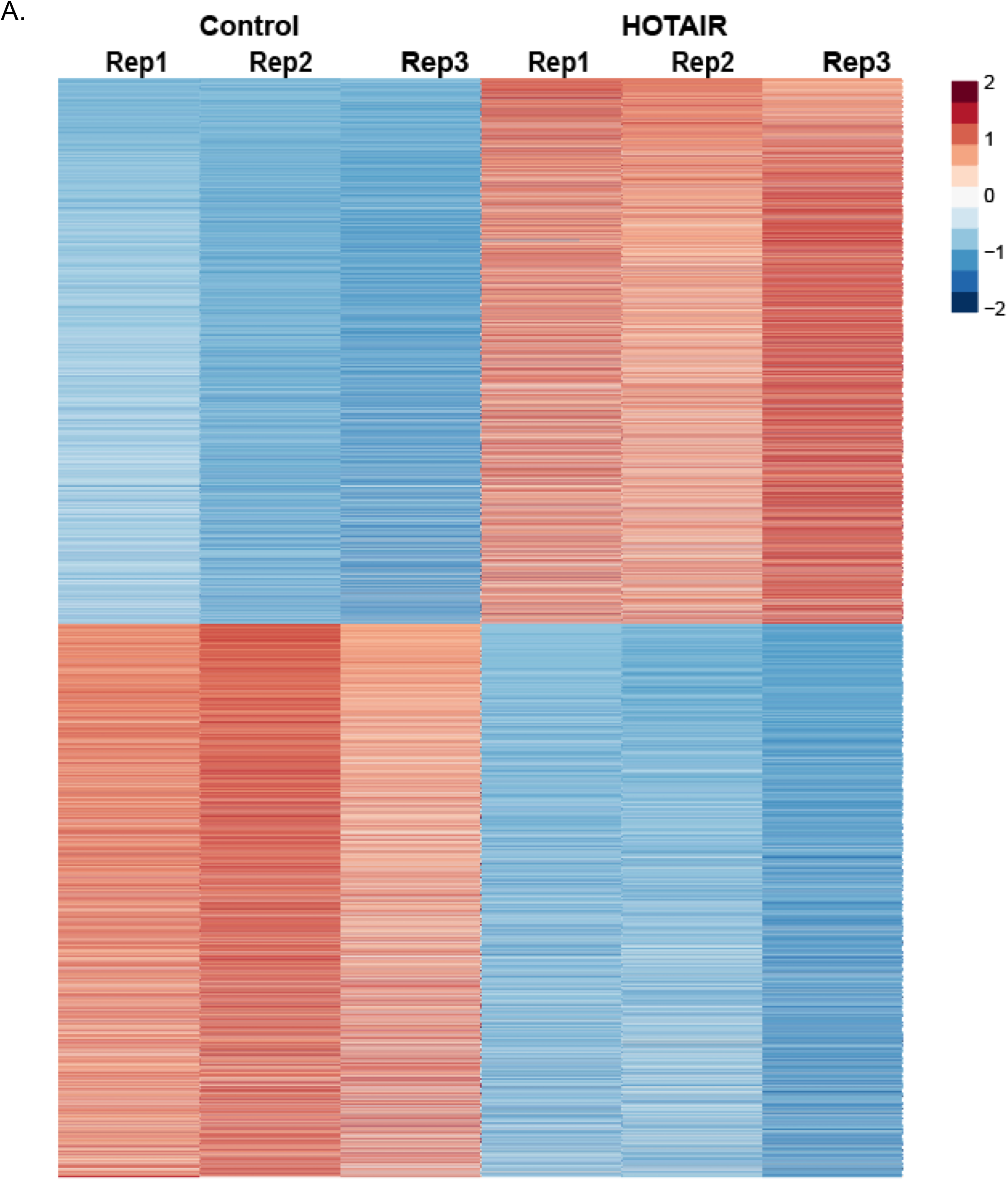
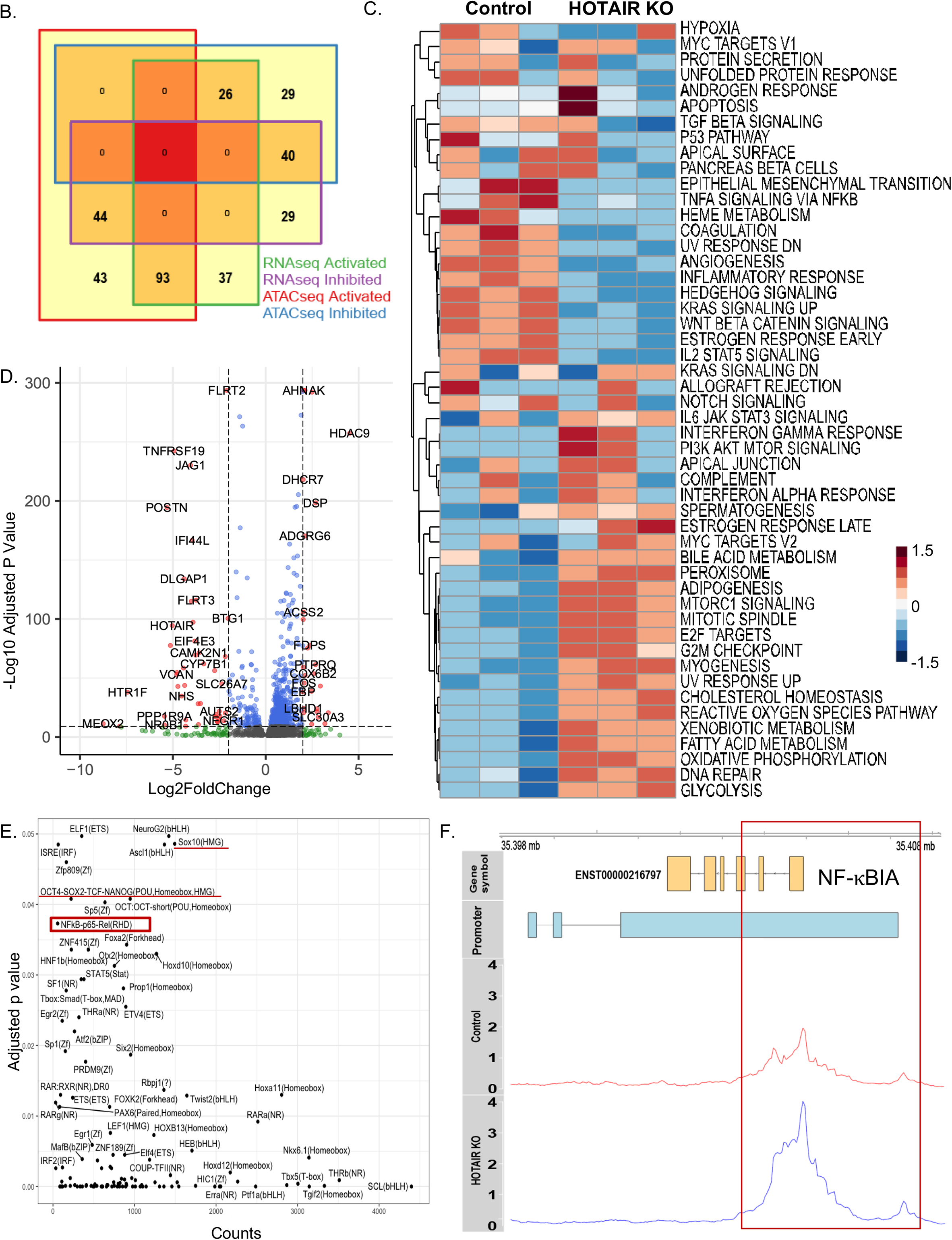
**A.** Heatmap of ATAC-seq total peaks for between HOTAIR KO vs. control Kuramochi clones. **B**. Venn diagram of activated and inhibited pathways from RNA-seq and ATAC-seq. IPA was utilized to analyze the activities of upstream regulators based on ATAC-seq and RNA-seq. **C.** Heatmap of activity scores for each biological process from the HALLMARK category of Molecular Signature Database between 2 groups**. D.** Volcano plot of the intersection of differentially expressed genes in RNA-seq and differential peaks-associated genes in ATAC-seq. **E.** Scatterplot of significantly (adjusted p value < 0.05) enriched motif based on peaks called to be differentially enriched in Control. **F.** Distribution of genome coverage from different samples in NFκBIA genomic region and the red box represents the region with differential peaks between two groups. ATAC-seq track analysis showed upregulated peaks in NFκBIA promoter in the HOTAIR KO cells compared to control cells.

**Supplemental Figure S4.**
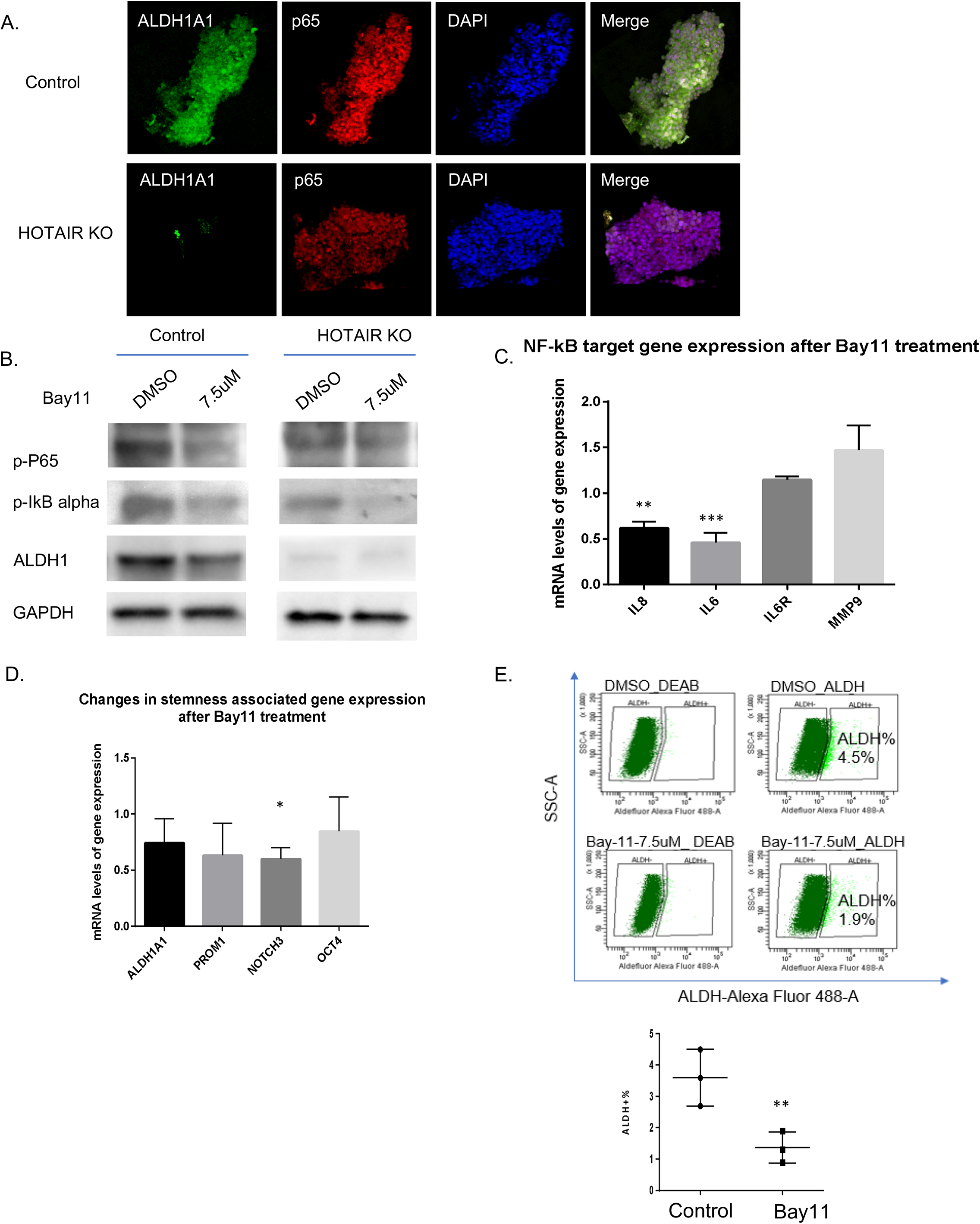
**A.** Spheroid staining of ALDH1A1 (green), p65 (red), and DAPI (blue) in Kuramochi HOTAIR KO clonal cells versus vector control cells. **B.** Western blot analysis p-P65 and p-IκB alpha and ALDH1 protein expression. GAPDH was used as loading control. **C.** NFκB target gene expression measured by quantitative real-time PCR (RT-qPCR) in control versus Bay- 11 treated cells. **D.** Stemness gene expression measured by RT-qPCR in control versus Bay-11 treated cells. **E.** Flow analysis (upper) and quantitation (lower) of ALDH+ population in control versus Bay-11 treated cells. *\*\*, P<0.01.* **F.** Scatter plot of ALDH+ population in control versus NF-κB KO Kuramochi cells.

**Supplemental Figure S5.**
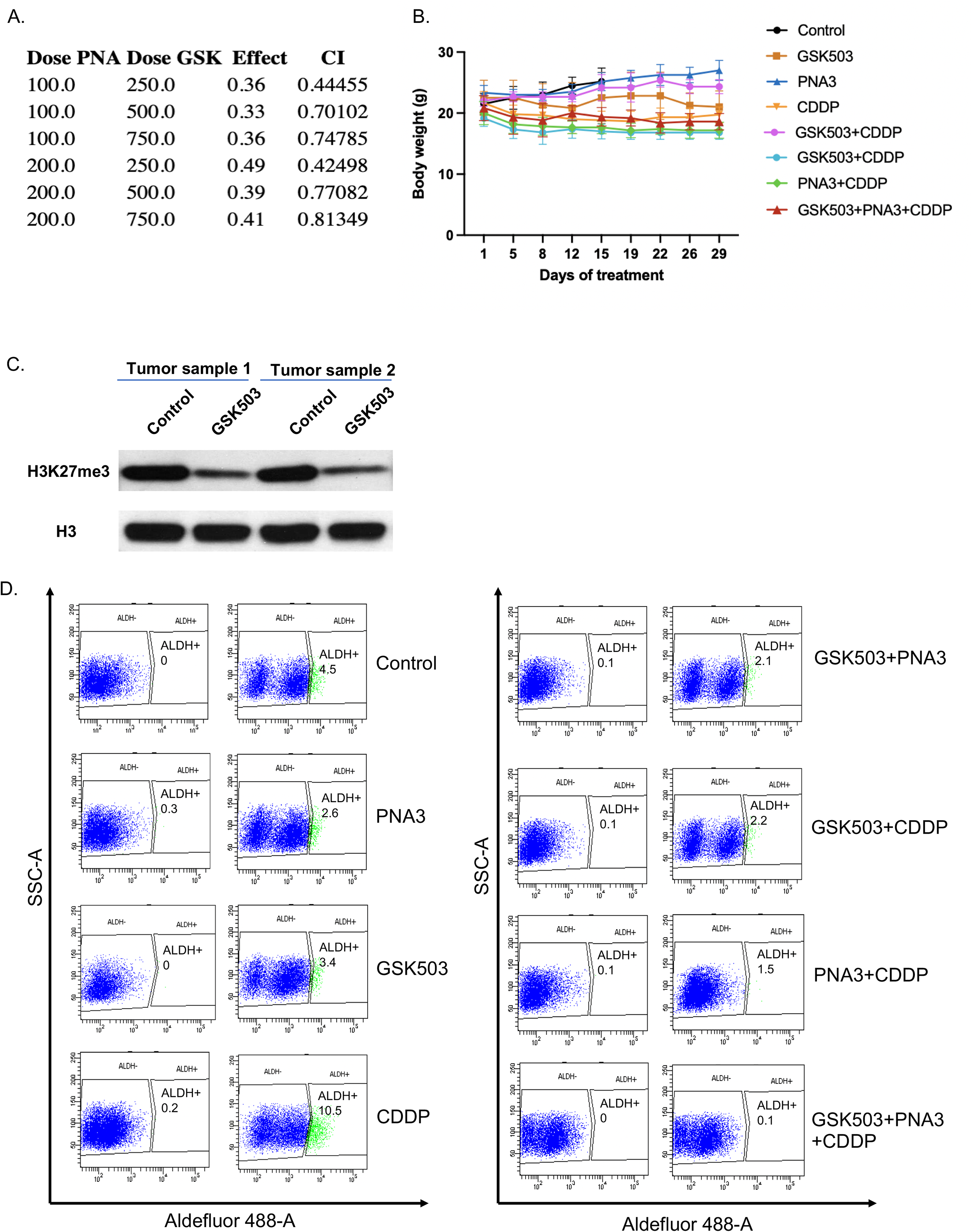
**A**. Synergistic inhibitory effects based on combination index value (CI value) lower than 1 were observed using a cell viability assay. **B.** Bodyweight of mice was recorded once the tumor size reached 200 mm^3^ in all 8 groups. **C.** H3K27me3 expression and loading control H3 expression in control vs.GSK503 treated mice were determined by western blot analysis. Tumor cells from 6 mice in each condition were grouped into two replicates. **D**. Flow data of dissociated tumor cells from 8 groups after treatments.

